# The neuronal plasticity gene *CRTC1* links survival with mitochondrial programs and tumor microenvironment features in Group 3 β medulloblastoma

**DOI:** 10.64898/2026.10.06.757146

**Authors:** Julia Vanini, Marina Müller Lupatini, André T. Brunetto, Caroline Brunetto de Farias, Mariane Jaeger, Rafael Roesler

## Abstract

CREB-regulated transcription coactivator 1 (CRTC1) regulates neuronal development, plasticity, and activity-dependent transcription, but its role in medulloblastoma (MB) remains poorly understood. We investigated expression of the *CRTC1* gene during human cerebellar development and across pediatric brain tumors and examined its molecular and clinical associations in MB using public transcriptomic datasets. *CRTC1* was broadly expressed in the developing cerebellum, with cell-state-specific variation within the granule-cell lineage. Among pediatric brain tumors, MB showed the highest median *CRTC1* expression, which differed significantly among MB molecular groups and subtypes, with highest levels in Group 4 MB. High *CRTC1* expression was associated with longer overall survival specifically in Group 3 β MB and remained significant after correction across the 12 molecular subtypes. Within Group 3 β tumors, *CRTC1* expression was inversely associated with enrichment of mitochondrial and bioenergetic processes, including oxidative phosphorylation, mitochondrial organization, and mitochondrial translation, whereas *CRTC1*-high and *CRTC1*-low tumors displayed distinct transcriptional profiles. *CRTC1* was also associated with selected cell-type-related transcriptional signatures related to the tumor microenvironment. These findings identify *CRTC1* as a marker of biologically and clinically relevant transcriptional heterogeneity in Group 3 β MB and support further investigation of its functional and prognostic significance.

**Graphical abstract:** 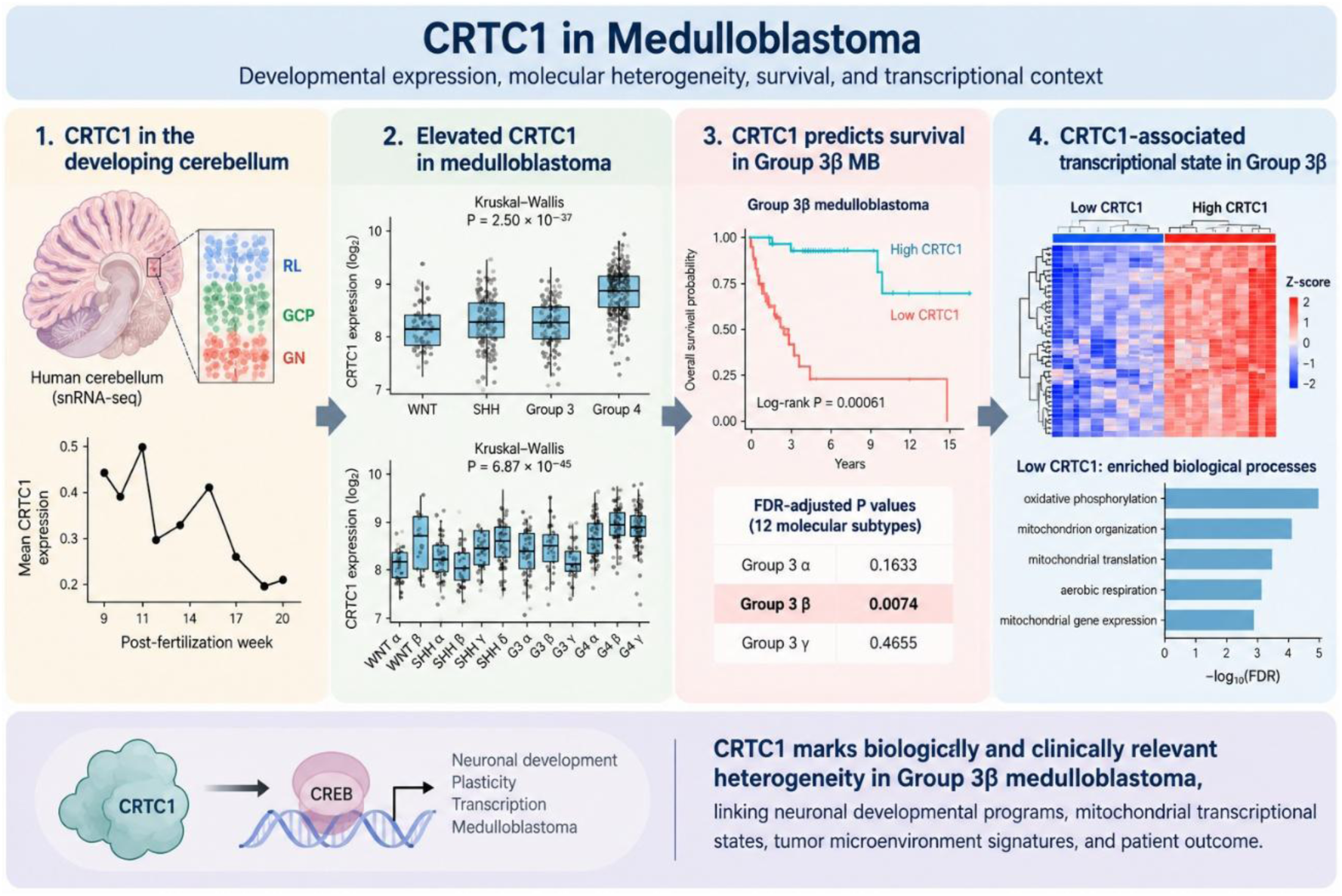

## 1 Introduction

Molecular mechanisms involved in neuronal development and synaptic plasticity may influence the biology of brain tumors [1, 2]. This emerging principle of cancer neuroscience is particularly relevant to medulloblastoma (MB), a malignant pediatric brain tumor arising in the developing cerebellum, in which developmental neural programs are intimately linked to tumor identity and molecular heterogeneity. MB comprises biologically distinct molecular groups and subtypes that reflect, at least in part, different developmental trajectories and cellular states [3–5]. Thus, molecules that normally couple neuronal signaling and plasticity to transcriptional responses may represent an underexplored component of MB biology.

The four defined molecular subgroups of MB are associated with distinct developmental origins in the hindbrain. Wingless (WNT) tumors are thought to arise from progenitors of the lower rhombic lip (RL)/dorsal brainstem, whereas Sonic Hedgehog (SHH) MBs originate predominantly from the granule neuron progenitor (GNP, also called granule cell precursor, GCP) lineage of the developing cerebellum. More recent developmental and single-cell studies have linked Group 3 and Group 4 tumors to the human rhombic lip and related glutamatergic lineages, with Group 3 displaying more primitive progenitor-like states and Group 4 showing greater resemblance to differentiating unipolar brush cell and glutamatergic neuronal lineages [3–7].

A refined molecular classification of MB emerged from the large-scale integrative analysis conducted by Cavalli et al. [3] within the Medulloblastoma Advanced Genomics International Consortium. Using transcriptomic and DNA methylation data from 763 primary tumors, the study resolved the four major molecular groups into 12 subtypes: two within Wingless (WNT), four within Sonic Hedgehog (SHH), and three each within Group 3 and Group 4. These subtypes display distinct genomic and clinical features, further emphasizing the biological heterogeneity of MB. Importantly, this level of molecular resolution provides a framework for identifying biological features and prognostic associations that may be confined to specific tumor subtypes and therefore obscured when analyses are restricted to the four molecular subgroups [8, 9].

In neurons, the CREB-regulated transcriptional coactivator (CRTC1, previously known as transducer of regulated CREB activity 1, TORC1) binds the transcription factor cAMP-response element-binding (CREB) to stimulate gene expression related to long-term synaptic plasticity [10, 11]. Synaptically localized CRTC1 in excitatory neurons translocates to the nucleus upon localized synaptic stimulation [12]. CRTC1 nuclear translocation in neurons in the lateral and basal nuclei of the amygdala accompanies fear memory retrieval and is required for memory reconsolidation [13]. Nuclear recruitment of CRTC1 in the basolateral amygdala accompanies formation of memory for contextual fear conditioning [14]. Impairment of CRTC1-dependent transcription by the β-amyloid peptide may be involved in memory impairment in Alzheimer’s disease (AD) models [15–17], and CRTC1 activates a transcriptional program which is deregulated at the early stages of AD [18]. Aberrant activation of CRTC1 is also involved in peripheral tumor types [19–23].

This evidence raises the possibility that CRTC1 may play a role in pediatric brain tumors, including MB. Here, we examined *CRTC1* gene expression across the molecular landscape of MB and investigated its associations with patient survival, tumor transcriptional programs, and microenvironment-related features, with particular attention to molecular subtype-specific effects.

## 2 Materials and methods

### 2.1 *CRTC1* expression in different brain areas

The levels of CRTC1 expression in number of mRNA transcripts per million (nTPM) in the human cerebral cortex, hippocampal formation, amygdala, basal ganglia, and cerebellum was visualized in The Human Protein Atlas (HPA) database (https://www.proteinatlas.org/ENSG00000105662-CRTC1/brain). No formal statistical analysis was applied.

### 2.2 Analysis of *CRTC1* expression in the developing human cerebellum

*CRTC1* expression during normal human cerebellar development was examined using two independent single-nucleus RNA-sequencing datasets, following an analytical framework previously applied [24]. In the dataset described by Aldinger et al. [25], comprising 69,174 nuclei from human cerebellar samples spanning 9–21 post-fertilization weeks (PFW), cell-type and developmental-stage annotations provided by the original study were retained. *CRTC1* expression was summarized across annotated cerebellar cell populations as mean expression and the percentage of nuclei with detectable expression. Expression was also examined descriptively across developmental stages. Because developmental stage was strongly associated with donor identity and cellular composition, developmental-stage profiles were not subjected to inferential trend testing.

To specifically examine the granule-cell developmental lineage, RL, GCP, and granule neuron (GN) populations were analyzed. Because nuclei derived from the same donor are not statistically independent, expression was first averaged within each cell state for each donor. Only donors represented in all three states were included in paired statistical analyses. Differences among RL, GCP, and GN states were evaluated using the Friedman test, followed by paired Wilcoxon signed-rank tests with Holm correction for multiple comparisons. Independent analysis was performed using the developmental human cerebellum dataset of Sepp et al. [26] (180,956 nuclei). Analysis was restricted to the RL/external granule layer (EGL) lineage and the developmental states GCP, GC_diff_1, GC_defined, and GC_diff_2. For each state, mean *CRTC1* expression and the percentage of nuclei with detectable expression were calculated using the original cell-state annotations. All statistical analyses were performed in R version 4.5.3 (R Foundation for Statistical Computing, Vienna, Austria) using RStudio version 2026.9.0.174 (Posit Software, PBC).

### 2.3 *CRTC1* expression across pediatric brain tumor types

To examine *CRTC1* expression across pediatric brain tumors, RNA-sequencing data were obtained from the Open Pediatric Brain Tumor Atlas (OpenPBTA) [27], release v23 (release-v23-20230115) (https://github.com/AlexsLemonade/OpenPBTA-analysis).

Gene-level expression values from the stranded RNA-seq by expectation-maximization (RSEM) nTPM matrix were matched to tumor annotations using Kids First Biospecimen identifiers. Analyses included the seven diagnostic groups represented by at least 30 samples: high-grade glioma/astrocytoma (HGG; *N* = 93), low-grade glioma/astrocytoma (LGG; *N* = 86), pilocytic astrocytoma (PA; *N* = 131), MB (*N* = 121), atypical teratoid/rhabdoid tumor (ATRT; *N* = 30), diffuse midline glioma (DMG; *N* = 34), and ependymoma (EPN; *N* = 93). Differences in *CRTC1* transcription among tumor types were evaluated using the Kruskal–Wallis test, followed by Dunn’s test with Benjamini– Hochberg (BH) correction for multiple comparisons. All statistical analyses were performed in R version 4.5.3 using RStudio version 2026.9.0.174.

### 2.4 *CRTC1* expression across MB molecular subgroups and subtypes

*CRTC1* expression was examined in the Cavalli MB cohort (GSE85217) [3], comprising 763 primary tumors classified into the four major molecular groups (WNT, SHH, Group 3, and Group 4) and 12 molecular subtypes (WNT α and β; SHH α, β, γ, and δ; Group 3 α, β, and γ; and Group 4 α, β, and γ). Expression values for *CRTC1* (ENSG00000105662) were extracted from the normalized gene-expression matrix and matched to the molecular annotations of each tumor. Differences in *CRTC1* transcription levels among the four molecular groups and among the 12 subtypes were evaluated separately using Kruskal– Wallis tests. When a significant overall difference was detected, pairwise comparisons were performed using Dunn’s post hoc test, with BH correction for multiple comparisons. Expression distributions were visualized using boxplots showing the median and interquartile range, with individual tumors displayed as jittered points. All statistical analyses were performed in R version 4.5.3 using RStudio version 2026.9.0.174.

### 2.5 *CRTC1* expression in MB single-cell RNA sequencing data

MB single-cell RNA sequencing (scRNA-seq) data from the dataset GSE156053 [28], available through the Gene Expression Omnibus (GEO) repository was used to visualize *CRTC1* expression at the single-cell level in MB. Data were explored using the Pediatric Neuro-oncology Cell Atlas interactive platform (https://www.pneuroonccellatlas.org/).

Uniform Manifold Approximation and Projection (UMAP) embeddings were used to visualize CRTC1 expression patterns across the different MB subgroups.

### 2.6 *CRTC1* survival associations in MB molecular subtypes

*CRTC1* expression and overall survival (OS) data from the Cavalli MB cohort [3] were analyzed separately within each of the 12 MB subtypes. Only tumors with available subtype assignment, *CRTC1* expression, OS time, and survival status were included. Within each subtype, the optimal *CRTC1* expression threshold was identified using the surv_cutpoint function of the R package survminer, with minprop = 0.20, requiring at least 20% of evaluable tumors in each expression category. Tumors were subsequently classified as low or high *CRTC1* using surv_categorize. OS was estimated by the Kaplan– Meier method and compared using the log-rank test. Univariable Cox proportional-hazards models were used to estimate hazard ratios (HRs) and 95% confidence intervals (CIs) for high versus low *CRTC1* expression. To account for testing across the 12 molecular subtypes, log-rank *P* values were adjusted using the BH procedure.

### 2.7 *CRTC1* coexpression analysis

To characterize the transcriptional context of *CRTC1* expression within Group 3 β MB, gene coexpression analysis was performed using expression data from all 37 Group 3 β tumors in the Cavalli cohort [3], irrespective of the availability of survival information. Spearman correlation coefficients were calculated between *CRTC1* expression and the expression of each gene represented in the dataset, and genes were ranked according to their correlation coefficient with *CRTC1*. For visualization of the coexpression structure, the 30 genes showing the strongest positive correlations with *CRTC1* were selected together with *CRTC1*. Expression values were standardized by gene (z-score), and hierarchical clustering was used to visualize coordinated expression patterns across tumors.

### 2.8 Gene Ontology enrichment

Functional enrichment analysis was performed separately for genes strongly positively or negatively correlated with *CRTC1*, defined as Spearman ρ ≥ 0.50 for positively correlated genes and ρ ≤ −0.50 for negatively correlated genes. This threshold identified 401 positively and 431 negatively correlated genes in Group 3 β MB. Gene symbols were converted to Entrez Gene IDs using the org.Hs.eg.db annotation database, resulting in 369 and 403 mapped genes, respectively. Gene Ontology (GO) Biological Process over-representation analysis was performed using the clusterProfiler R package. All genes from the Cavalli expression dataset successfully mapped to Entrez Gene IDs (*N* = 18,930) were used as the background universe. Enrichment *P* values were adjusted for multiple testing using the BH method, and GO terms with adjusted *P* < 0.05 were considered significant.

### 2.9 Differential gene expression according to *CRTC1* expression in Group 3 β MB

Differential gene expression was investigated in Group 3 β MB (*N* = 37) from the Cavalli cohort [3], using the GlioVis data portal (https://gliovis.bioinfo.cnio.es/) [29]. Tumors were stratified according to *CRTC1* expression, and gene expression profiles were compared between *CRTC1*-high and *CRTC1*-low tumors using the GlioVis “High versus Low” analysis. Genes displaying differential expression between the two groups were visualized as a hierarchically clustered heatmap. Relative expression levels are represented by the heatmap color scale.

### 2.10 Tumor microenvironment analysis

Tumor microenvironment-associated transcriptional features were investigated in Group 3 β MB tumors with available expression data (*N* = 37) from the Cavalli cohort [3]. Gene expression data were converted from Ensembl identifiers to HUGO Gene Nomenclature Committee (HGNC) gene symbols using the org.Hs.eg.db annotation package. Ambiguous mappings and duplicated gene symbols were removed, resulting in a gene-level expression matrix containing 19,821 unique gene symbols. Sample identities and ordering were verified against the clinical and molecular annotation data before downstream analyses. Immune and stromal components were first evaluated using ESTIMATE (R package version 1.0.13). Expression data were filtered against the ESTIMATE common-gene set, resulting in 9,422 genes available for scoring. StromalScore, ImmuneScore, ESTIMATEScore, and inferred tumor purity were calculated for each tumor using the Affymetrix platform setting. Associations between *CRTC1* expression and ESTIMATE-derived measures were assessed using Spearman rank correlation and linear regression models using standardized *CRTC1* expression. BH correction was applied across the four ESTIMATE-derived outcomes.

Cell-type-associated transcriptional enrichment was subsequently examined using xCell (version 1.1.0) within the 37 Group 3 β MB tumors. Associations between CRTC1 expression and each of the 64 xCell cell-type enrichment scores were evaluated using Spearman rank correlation, followed by BH correction across the 64 signatures. Aggregate xCell ImmuneScore, StromaScore, and MicroenvironmentScore were evaluated separately. xCell analysis was performed using the 19,821-gene expression matrix. The analysis generated enrichment scores for 64 cell-type signatures, together with aggregate ImmuneScore, StromaScore, and MicroenvironmentScore measures. xCell scores were interpreted as gene-expression-based enrichment signatures rather than direct estimates of cellular fractions or absolute cell abundance.

## 3 Results

### 3.1 *CRTC1* is dynamically expressed during human cerebellar development

MB occurs in the cerebellum and all the molecular subgroups and subtypes of MB are thought to arise from distinct cell populations and developmental trajectories within the developing hindbrain, particularly the cerebellum and RL, highlighting the relevance of examining candidate genes in their normal developmental context. Studies investigating the function of CRTC1 in the adult brain have focused on areas including the hippocampus and basolateral amygdala, whereas its role in the cerebellum remains largely unexplored. Data from The Human Protein Atlas show high mRNA expression of *CRTC1* in the adult cerebellum compared to other brain areas (Fig. 1). We found that the *CRTC1* gene was broadly detectable across cell populations of the developing human cerebellum in the Aldinger dataset, although both mean expression and the proportion of *CRTC1*-positive nuclei varied substantially among cell types (Fig. 2A). The highest mean expression was observed in endothelial cells (mean = 0.986; 46.6% positive), followed by brainstem choroid/ependymal cells (0.665; 38.9%) and pericytes (0.590; 34.4%). Within the granule-cell lineage, aggregate mean expression was similar in RL (0.343; 22.8% positive) and GCP (0.345; 21.0%) populations and lower in GN (0.221; 15.6%).

**Fig. 1.**
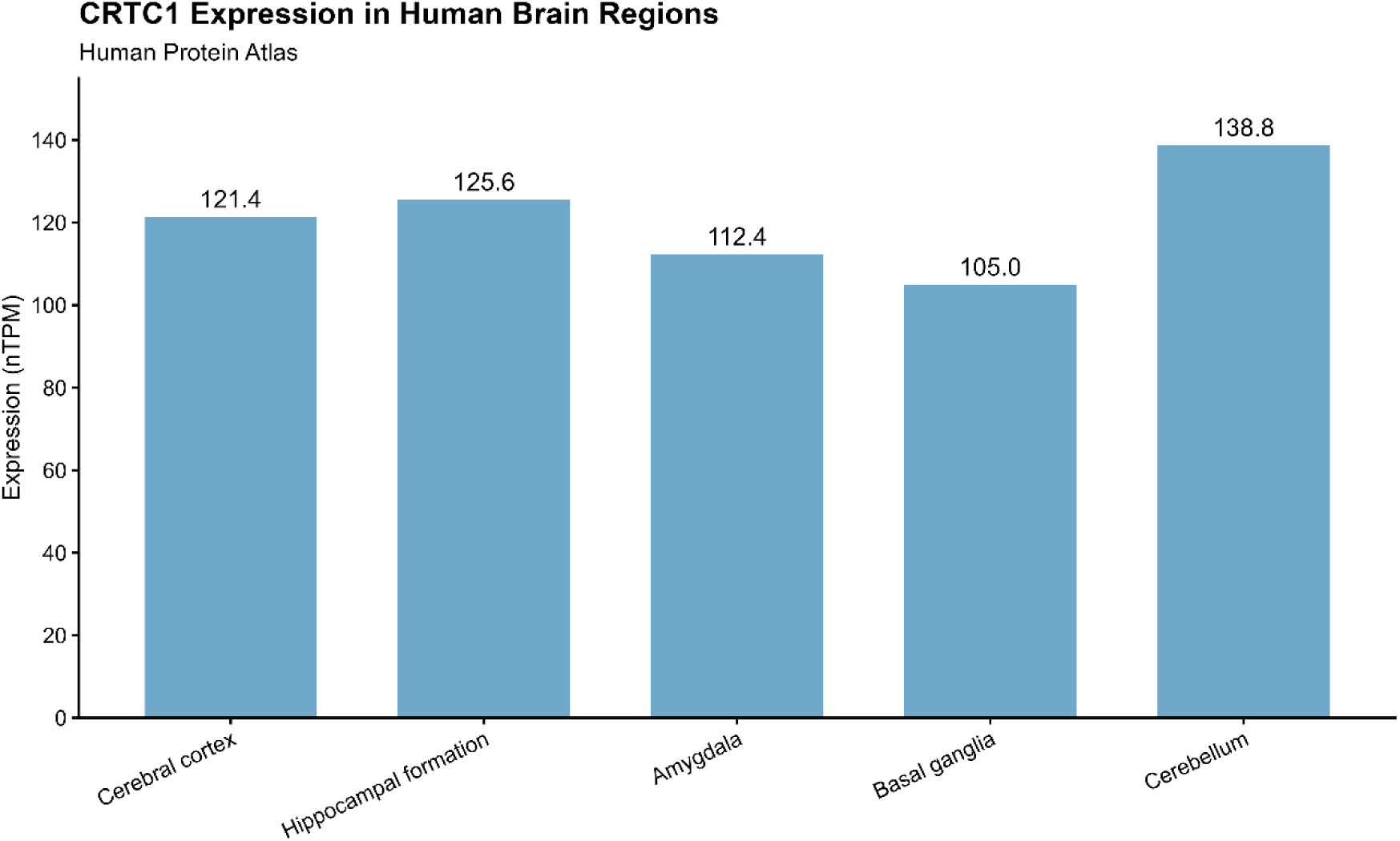
Messenger RNA expression of *CRTC1* in the human cerebral cortex, hippocampal formation, amygdala, basal ganglia, and cerebellum. Data normalized as nTPM were obtained from The Human Protein Atlas (https://www.proteinatlas.org/).

**Fig. 2.**
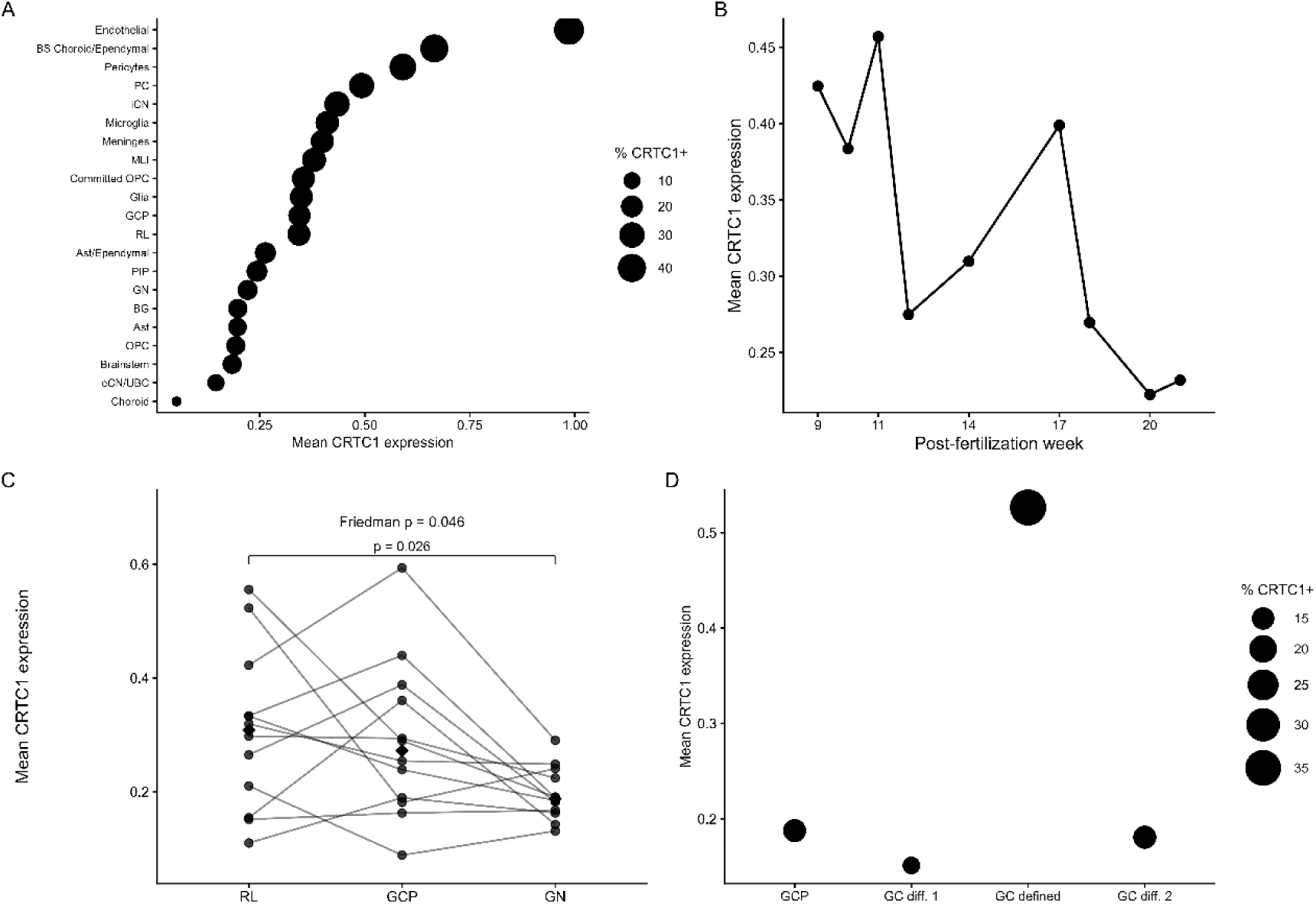
C*R*TC1 expression during human cerebellar development. **A** *CRTC1* expression across annotated cell populations in the Aldinger et al. developing human cerebellum single-nucleus RNA-sequencing dataset (69,174 nuclei) (Aldinger et a. 2021). Dot position represents mean *CRTC1* expression and dot size represents the percentage of nuclei with detectable *CRTC1* expression. **B** Mean *CRTC1* expression across developmental stages from 9 to 21 PFW. Developmental stage values are presented descriptively because developmental age is associated with donor identity and cell-type composition in this dataset. **C** Donor-level mean *CRTC1* expression across RL, granule cell precursor GCP, and GN populations. Lines connect populations derived from the same donor; diamonds indicate median values. Only donors represented in all three states were included (*N* = 11). Differences were evaluated using the Friedman test (*p* = 0.046), followed by paired Wilcoxon signed-rank tests with Holm correction. *CRTC1* expression was lower in GN than RL (adjusted *p* = 0.026). **D** Independent analysis of the Sepp et al. [26] RL/EGL lineage showing mean *CRTC1* expression across GCP, GC_diff_1, GC_defined, and GC_diff_2 states. Dot size represents the percentage of *CRTC1*-positive nuclei. The GC_defined population showed markedly higher *CRTC1* expression and detection frequency than the other analyzed states. Ast, astrocytes; Ast/Ependymal, astrocyte/ependymal cells; BG, basal ganglia; BS Choroid/Ependymal, brainstem choroid/ependymal cells; eCN/UBC, excitatory cerebellar nuclei/unipolar brush cells; GC, granule cell; GC defined, defined granule cells; GC diff. 1, granule cell differentiation state 1; GC diff. 2, granule cell differentiation state 2; GCP, granule cell progenitors; GN, granule neurons; iCN, inhibitory cerebellar nuclei; MLI, molecular layer interneurons; OPC, oligodendrocyte precursor cells; PC, Purkinje cells; PIP, Purkinje cell/interneuron progenitors; RL, rhombic lip.

Across developmental age, *CRTC1* remained detectable throughout the 9–21 PFW interval, with variation in mean expression across developmental stages (Fig. 2B). Mean expression ranged from 0.222 at 20 PFW to 0.457 at 11 PFW. These profiles were considered descriptive because most developmental stages were represented by only one or a small number of donors and differed in cellular composition.

Donor-level analysis of the RL–GCP–GN lineage included 11 donors represented in all three developmental states (Fig. 2C). Mean *CRTC1* expression differed across the three populations (Friedman χ² = 6.17, df = 2, *P* = 0.0458). Pairwise analysis showed lower expression in GN relative to RL (Holm-adjusted *P* = 0.0258). The GCP–GN comparison showed a similar direction but did not remain significant after correction (raw *P* = 0.0310; Holm-adjusted *P* = 0.0620), whereas RL and GCP did not differ (*P* = 1.00).

Analysis of the independent Sepp dataset revealed a distinct pattern of developmental-state-specific *CRTC1* expression within the RL/ EGL lineage (Fig. 2D). Mean expression and detection frequency were relatively similar in GCP (mean = 0.188; 15.3% positive), granule cell (GC)_diff_1 (0.151; 13.5%), and GC_diff_2 (0.181; 15.5%). In contrast, the GC_defined population showed markedly higher *CRTC1* expression (mean = 0.526; 36.0% positive). Further subdivision of the differentiating populations into early and late states did not account for this difference: mean expression ranged from 0.145 to 0.192 across GC_diff_1_early, GC_diff_1_late, GC_diff_2_early, and GC_diff_2_late, indicating that the elevated signal was specifically associated with the GC_defined state.

### 3.2 *CRTC1* is highly expressed in MB relative to other pediatric brain tumors

*CRTC1* expression differed significantly among the seven pediatric brain tumor types (Kruskal–Wallis χ^2^ = 62.5, df = 6, *P* = 1.40 × 10^−11^; Fig. 3). MB showed the highest median *CRTC1* expression (11.0 nTPM), compared with EPN (8.41), LGG (7.24), PA (6.78), HGG (6.45), DMG (6.04), and ATRT (4.68 nTPM). Pairwise analyses confirmed significantly higher *CRTC1* expression in MB than in each of the other six tumor groups (BH-adjusted Wilcoxon *P* = 1.9 × 10^−6^ vs HGG, 0.00058 vs LGG, 2.0 × ^−8^ vs PA, 2.7 × 10^−5^ vs ATRT, 9.5 × 10^−5^ vs DMG, and 0.00468 vs EPN). Thus, among the major pediatric brain tumor types examined, MB was characterized by comparatively high *CRTC1* expression.

**Fig. 3.**
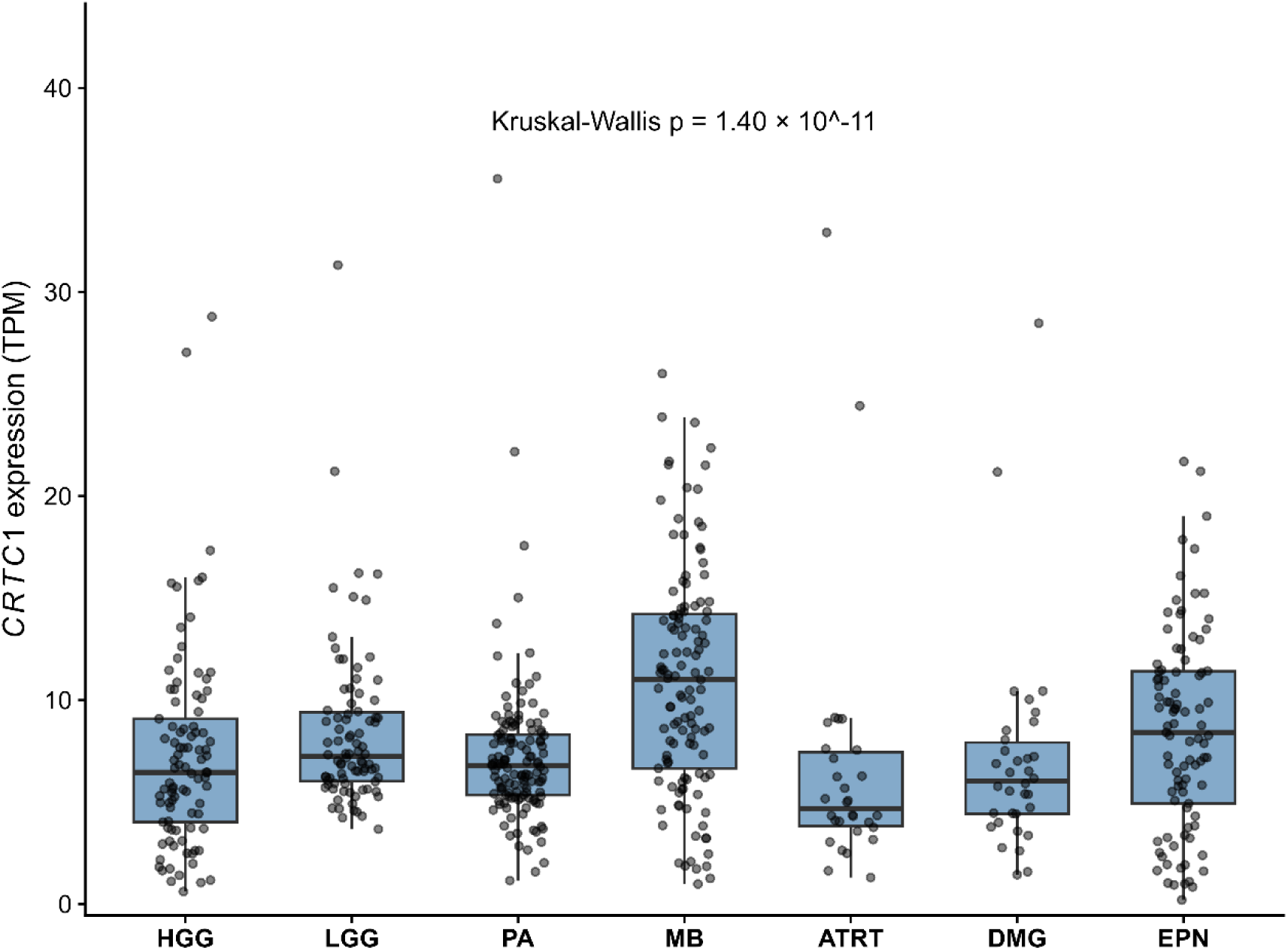
C*R*TC1 expression across pediatric brain tumor types. Gene expression was examined in high-grade glioma/astrocytoma (HGG; *N* = 93), low-grade glioma/astrocytoma (LGG; *N* = 86), pilocytic astrocytoma (PA; *N* = 131), medulloblastoma (MB; *N* = 121), atypical teratoid/rhabdoid tumor (ATRT; *N* = 30), diffuse midline glioma (DMG; *N* = 34), and ependymoma (EPN; *N* = 93). Expression is shown as nTPM. Boxes represent the median and interquartile range, and points represent individual tumor samples. Differences among groups were assessed using the Kruskal– Wallis test (*P* = 1.40 × 10^−11^), followed by pairwise Wilcoxon rank-sum tests with BH correction.

### 3.3 *CRTC1* expression differs across MB molecular subgroups and subtypes

*CRTC1* was expressed across all four major molecular groups and all 12 molecular subtypes of MB, but its expression differed significantly according to molecular classification. At the molecular subgroup level, *CRTC1* expression differed significantly among WNT, SHH, Group 3, and Group 4 tumors (Kruskal–Wallis χ^2^ = 173.27, df = 3, *P* < 2.2 × 10^−16^; Fig. 4A). Post hoc Dunn tests with BH correction showed that Group 4 tumors had significantly higher *CRTC1* expression than WNT (adjusted *P* = 4.52 × 10^−17^), SHH (adjusted *P* = 5.81 × 10^−24^), and Group 3 tumors (adjusted *P* = 5.88 × 10^−22^). In contrast, *CRTC1* expression did not differ significantly among WNT, SHH, and Group 3 tumors (all adjusted *P* ≥ 0.15).

**Fig. 4.**
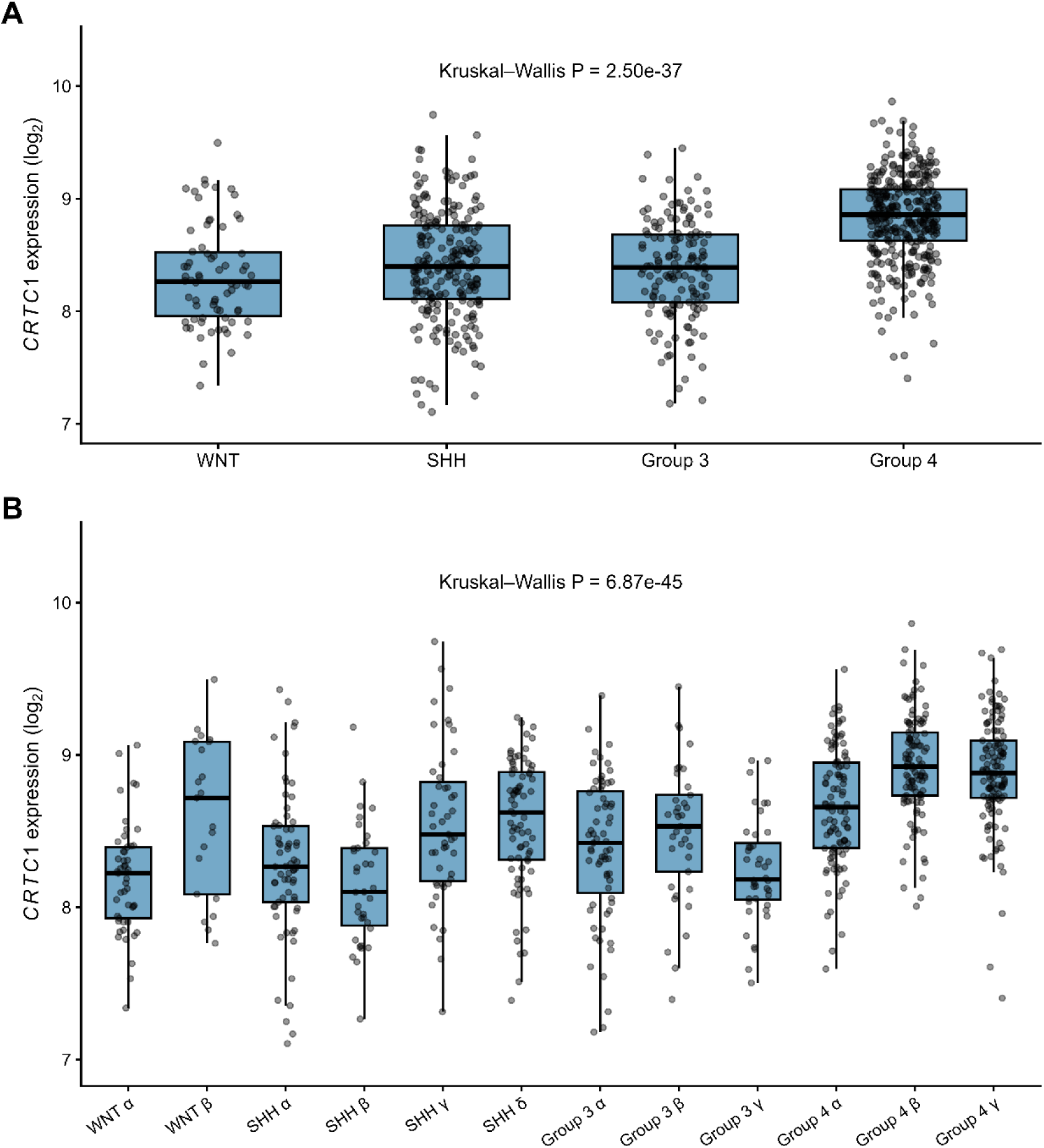
C*R*TC1 expression across molecular groups and subtypes of MB. Transcription levels were examined in 763 primary MB tumors from the Cavalli cohort (GSE85217) [3]. **A** Distribution of *CRTC1* expression across the four major molecular groups: WNT, SHH, Group 3, and Group 4. *CRTC1* expression differed significantly among groups (Kruskal–Wallis *P* = 2.50 × 10^−45^). **B** Distribution of *CRTC1* expression across the 12 molecular subtypes. Expression differed significantly among subtypes (Kruskal–Wallis *P* = 6.87 × 10^−37^). Boxes indicate the interquartile range with the median shown by the central line; individual points represent individual tumors. Pairwise comparisons were performed using Dunn’s post hoc test with BH correction for multiple testing. Group 4 tumors expressed significantly higher *CRTC1* levels than each of the other molecular subgroups. At the subtype level, significant within-group differences included WNT β versus WNT α, SHH γ/δ versus SHH α/β, Group 3 β versus Group 3 γ, and Group 4 β/γ versus Group 4 α.

Significant heterogeneity was also observed across the 12 molecular subtypes (Kruskal–Wallis χ^2^ = 238.57, df = 11, P < 2.2 × 10^−16^; Fig. 4B). Within WNT MB, *CRTC1* expression was significantly higher in WNT β than WNT α tumors (adjusted *P* = 0.0013). Within SHH MB, SHH γ and SHH δ showed higher expression than SHH α (adjusted *P*= 0.0166 and 0.0013, respectively) and SHH β (adjusted *P* = 9.96 × 10^−4^ and 5.33 × 10^−5^, respectively), whereas SHH γ and SHH δ did not differ from each other (adjusted *P* = 0.699). Within Group 3, Group 3 β showed significantly higher *CRTC1* expression than Group 3 γ (adjusted *P* = 0.0195), whereas Group 3 α did not differ significantly from either Group 3 β or Group 3 γ. Within Group 4, expression was higher in both Group 4 β and Group 4 γ than in Group 4 α (adjusted *P* = 5.46 × 10^−5^ and 5.05 × 10^−5^, respectively), while Group 4 β and Group 4 γ did not differ significantly (adjusted *P* = 0.578). Thus, *CRTC1* expression is characterized by both a marked elevation in Group 4 MB overall and significant heterogeneity among molecular subtypes within several major groups. Single-cell RNA sequencing data further allows visualization of CRTC1 expression across MB subgroups (Fig. 5).

**Fig. 5.**
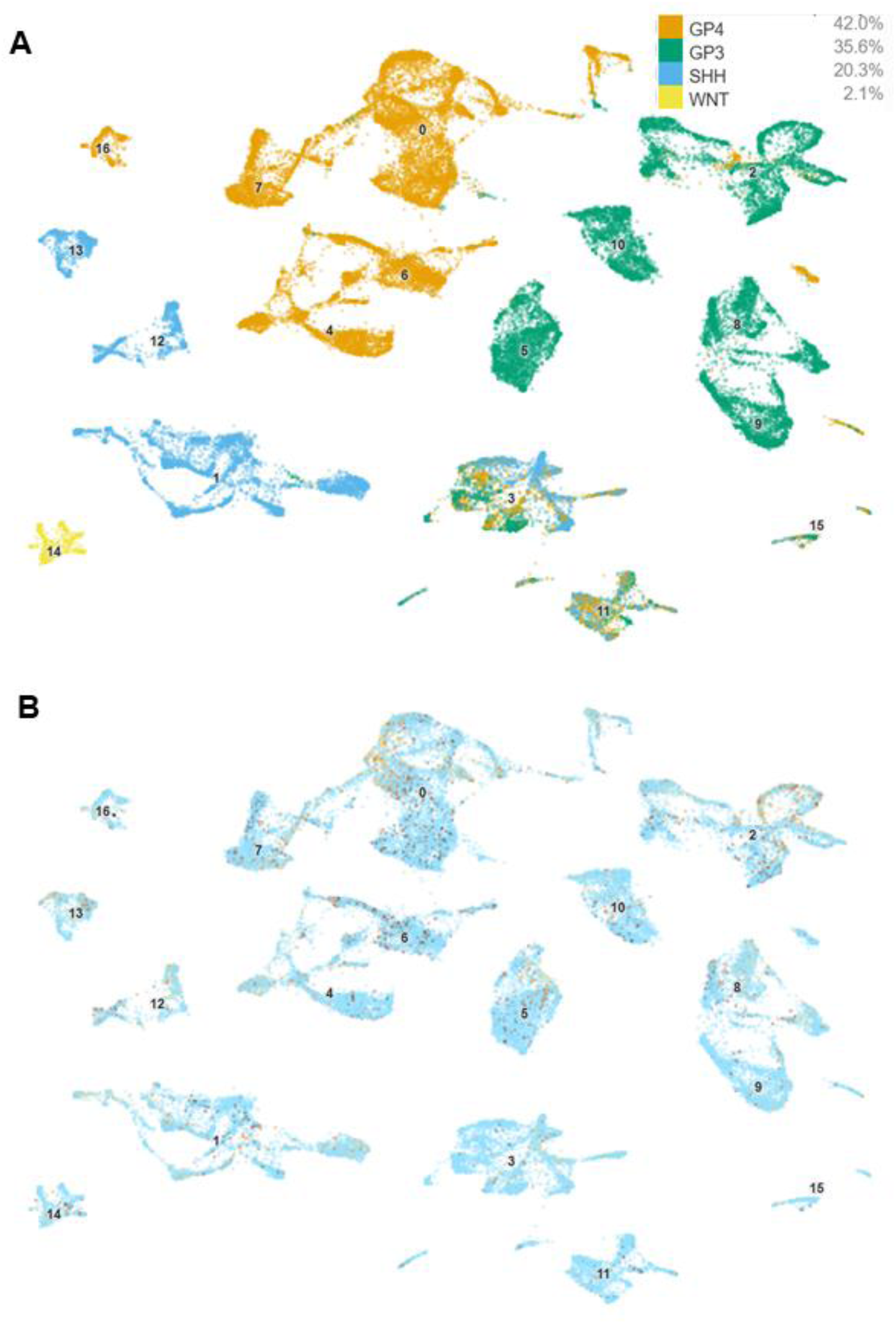
UMAP plots of expression of *CRTC1* in the four MB subgroups at single cell resolution. **A** Cells annotated by molecular subgroup across 28 MB tumor samples. **B** Cells annotated by *CRTC1*. Data were accessed using single-cell scRNA-seq data available in GSE156053 [28] through the Pediatric Neuro-oncology Cell Atlas (https://www.pneuroonccellatlas.org/). G3, Group 3 MB; G4, Group 4 MB.

### 3.4 High *CRTC1* expression is associated with better survival selectively in Group 3 β MB

Among the 12 molecular subtypes, the only significant survival association was observed in Group 3 β MB (Fig. 6). Twenty-seven Group 3 β tumors had evaluable survival data, including 12 deaths. The maximally selected *CRTC1* cutoff was 8.500507, classifying 12 tumors as low and 15 as high *CRTC1*. High expression was associated with markedly longer OS (HR = 0.109, 95% CI 0.024–0.502; Cox *P* = 0.0045). The survival curves were strongly separated by the log-rank test (*P* = 0.00061), and the association remained significant after BH correction across all 12 subtype analyses (BH False Discovery Rate, FDR = 0.0074).

**Fig. 6.**
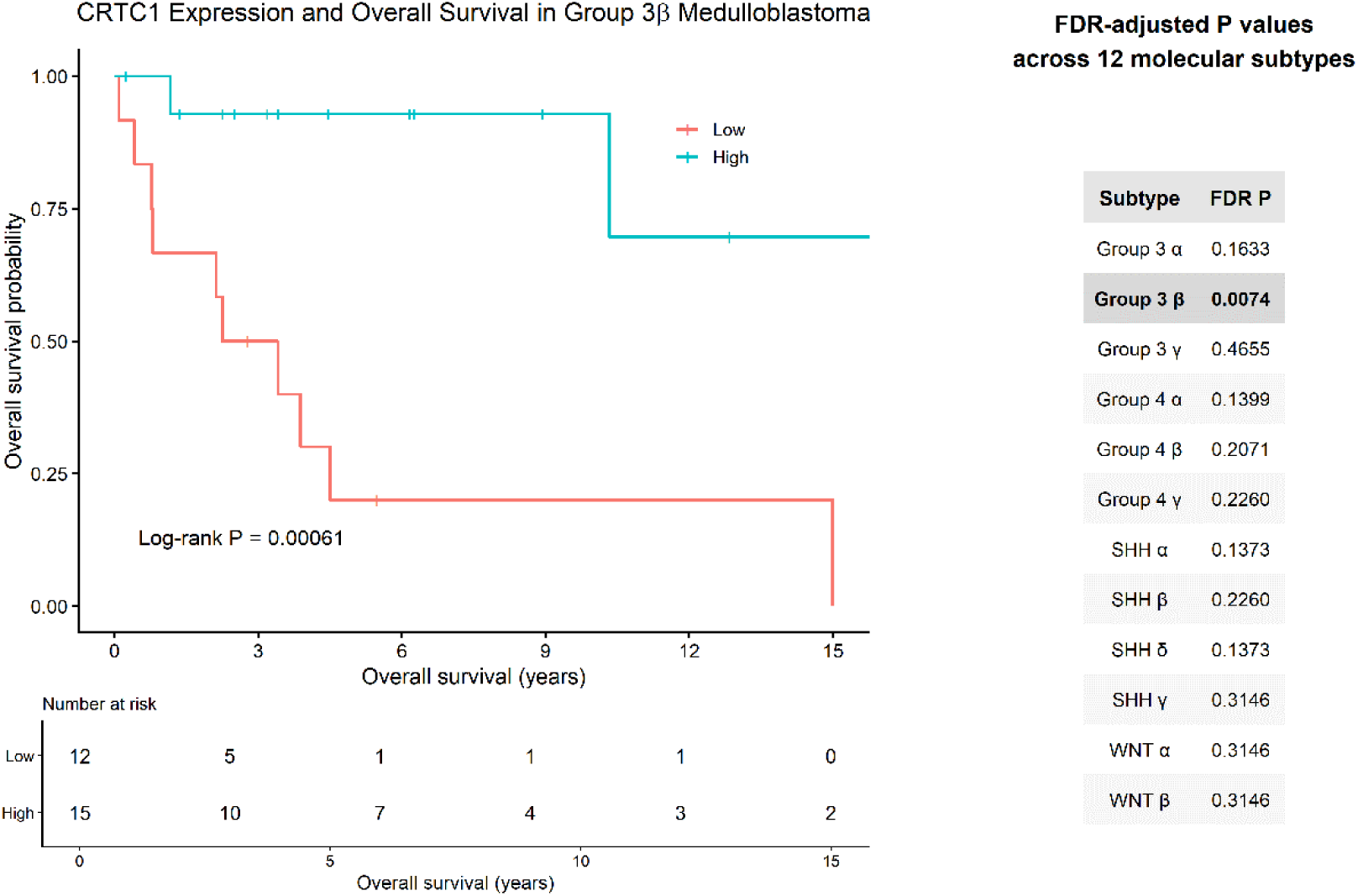
C*R*TC1 expression and OS in Group 3 β MB. Kaplan–Meier curves show OS according to low and high *CRTC1* expression in Group 3 β tumors from the Cavalli cohort (GSE85217) [3] (*N* = 27; 12 low and 15 high). Expression groups were defined using the maximally selected cutoff identified with survminer (cutoff = 8.500507; minprop = 0.20). High *CRTC1* expression was associated with longer OS (HR = 0.109, 95% CI 0.024– 0.502; log-rank *P* = 0.00061). The accompanying table shows BH-adjusted *P* values for survival analyses across all 12 molecular subtypes; Group 3 β was the only subtype remaining significant after multiple-testing correction (BH FDR = 0.0074). Tick marks indicate censored observations, and numbers at risk are shown below the Kaplan–Meier plot.

No other molecular subtype remained significant after multiple-testing correction. Nominal log-rank associations were observed in Group 4 α (*P* = 0.0466), SHH α (*P* = 0.0276), and SHH δ (*P* = 0.0343), but their corresponding BH FDR values were 0.1399, 0.1373, and 0.1373, respectively. Group 3 α showed a nonsignificant trend toward improved survival with high *CRTC1* (HR = 0.43; log-rank *P* = 0.0680; BH FDR = 0.1633). The WNT subtype estimates were unstable because very few deaths occurred (one event in WNTα and two in WNTβ), and their Cox estimates should therefore not be interpreted as reliable effect-size estimates.

### 3.5 *CRTC1* coexpression structure in Group 3 β MB reveals an inverse association with mitochondrial and bioenergetic programs

Given the specific association of Group 3 β MB with survival, we selected this subtype for further analyses. To investigate the biological context of *CRTC1* expression within Group 3 β MB, we examined gene coexpression across all 37 Group 3 β tumors. Strong positive correlations were observed between *CRTC1* and several genes, including *SPPL2B* (ρ = 0.810), *MAU2* (ρ = 0.781), *MED25* (ρ = 0.763), *TBC1D17* (ρ = 0.751), *WBP2* (ρ = 0.749), *SCAMP4* (ρ = 0.748), and *NUMBL* (ρ = 0.748). Hierarchical clustering of *CRTC1* and its 30 most strongly positively correlated genes demonstrated coordinated expression patterns across Group 3 β tumors (Fig. 7).

**Fig. 7.**
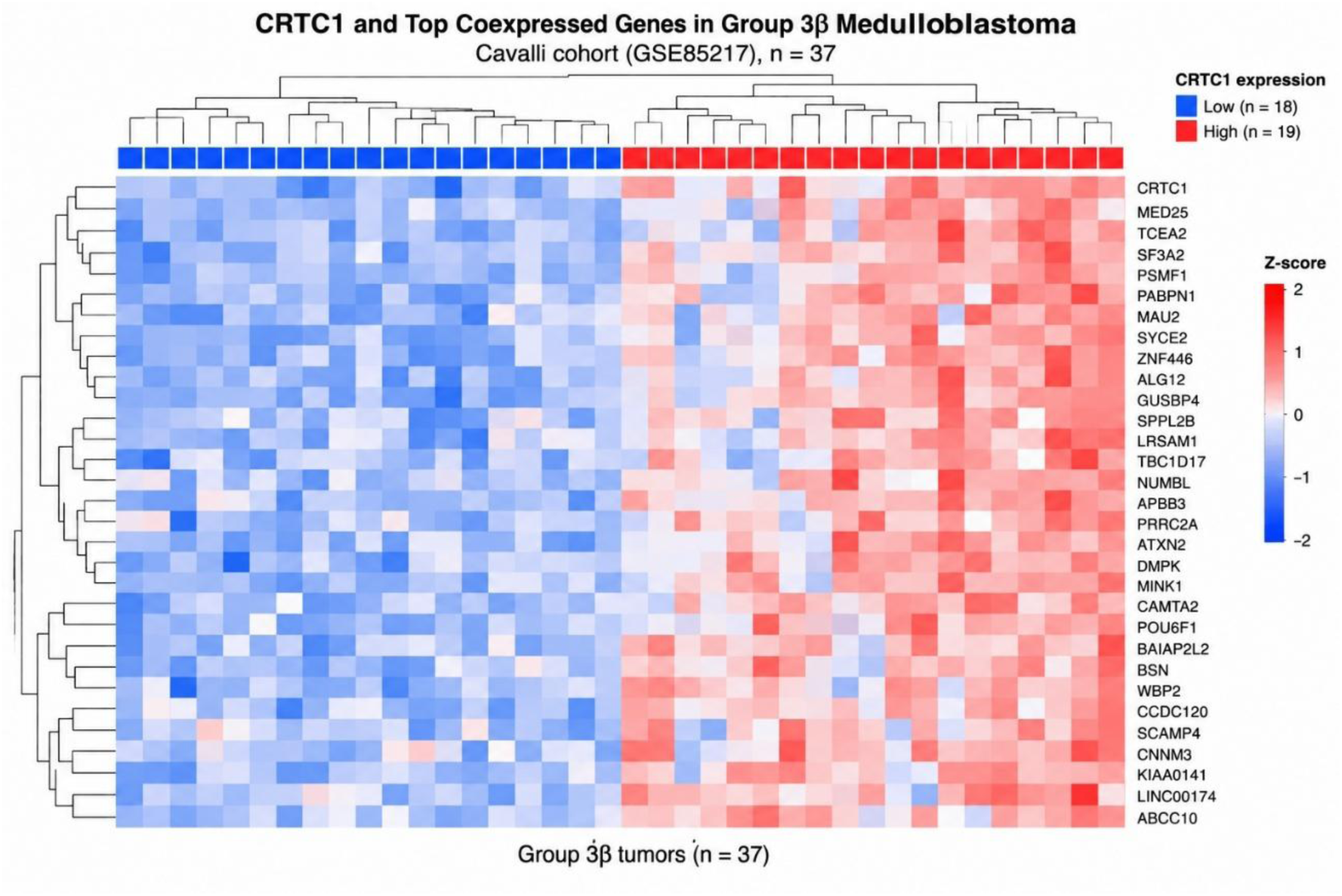
C*R*TC1 coexpression landscape in Group 3 β MB. Heatmap showing *CRTC1* and the 30 genes most strongly positively correlated with *CRTC1* expression across 37 Group 3 β tumors from the Cavalli cohort (GSE85217) [3]. Spearman correlation coefficients were calculated using expression data from all Group 3 β tumors, irrespective of survival data availability. Expression values were standardized by gene (z-scores), and hierarchical clustering was used to visualize coordinated expression patterns across tumors and genes.

GO Biological Process analysis in Group 3 β MB tumors revealed a marked functional asymmetry between genes positively and negatively correlated with *CRTC1*. Positively correlated genes showed relatively limited functional enrichment, with six significant GO terms, including processes related to transcriptional elongation and neural tube development. In contrast, genes negatively correlated with *CRTC1* showed strong and coherent enrichment for mitochondrial and bioenergetic processes. Among the most significantly enriched terms were mitochondrion organization (30 genes; adjusted *P* = 1.21 × 10^−5^), oxidative phosphorylation (16 genes; adjusted *P* = 1.21 × 10^−5^), ATP synthesis coupled electron transport (14 genes; adjusted *P* = 1.21 × 10^−5^), aerobic electron transport chain (13 genes; adjusted *P* = 2.26 × 10^−5^), mitochondrial translation (15 genes; adjusted *P* = 7.32 × 10^−5^), mitochondrial respiratory chain complex assembly (11 genes; adjusted *P* = 6.02 × 10^−4^), and mitochondrial gene expression (15 genes; adjusted *P* = 1.00 × 10^−3^) (Fig. 8). Additional significantly enriched processes included respiratory electron transport, aerobic respiration, cellular respiration, and ATP biosynthesis.

**Fig. 8.**
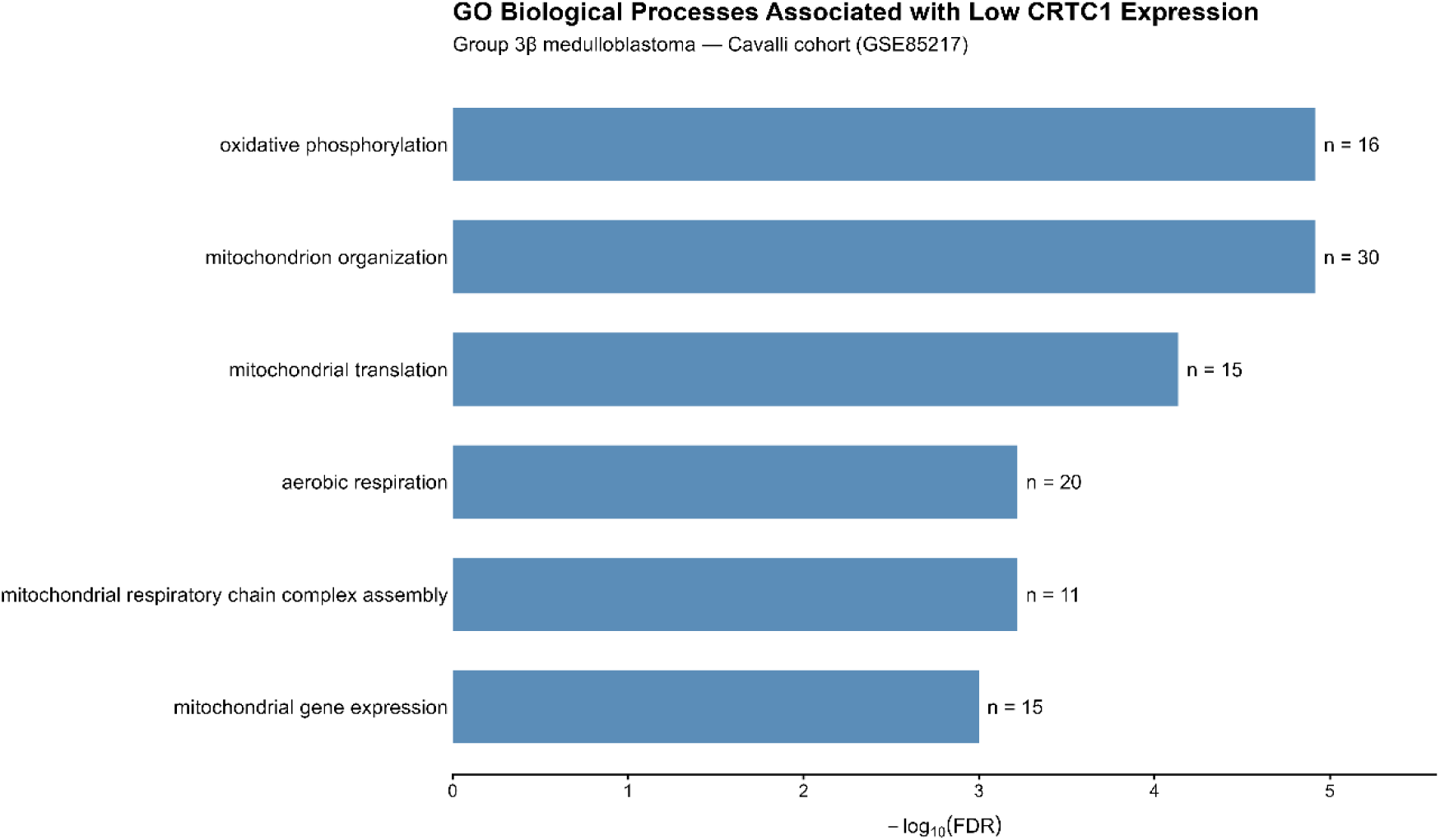
Gene Ontology (GO) Biological Process enrichment among genes negatively correlated with *CRTC1* in Group 3 β MB. Representative significantly enriched GO Biological Process terms among genes strongly negatively correlated with *CRTC1* expression (Spearman ρ ≤ −0.50) across 37 Group 3 β tumors from the Cavalli cohort (GSE85217) [3]. GO over-representation analysis was performed using genes represented in the expression dataset as the background universe, and *P* values were adjusted for multiple testing using the BH method. Bars represent −log10(FDR), and labels indicate the number of negatively correlated genes contributing to each GO term. Enriched processes predominantly involved mitochondrial organization, oxidative phosphorylation, mitochondrial translation and gene expression, respiratory chain complex assembly, and aerobic respiration.

Together, these findings indicate that lower *CRTC1* expression within Group 3 β MB is associated with increased expression of genes involved in mitochondrial organization, oxidative phosphorylation, respiratory electron transport, ATP production, and mitochondrial gene expression and translation.

### 3.6 *CRTC1* expression Is associated with distinct transcriptional profiles in Group 3 β MB

Comparison of *CRTC1*-high and *CRTC1*-low Group 3 β tumors revealed marked differences in gene expression, with hierarchical clustering identifying two broadly reciprocal transcriptional patterns (Fig. 9). Genes showing higher expression in association with higher *CRTC1* included *PCDH9*, *PDE6H*, *KCNJ5*, *GALNT12*, *PIEZO1*, *LINC00174*, *CCNJL*, *NFIX*, and *TCEA2*, among others. Conversely, genes including *GNB4*, *MRPS7*, *DCTPP1*, *MSRB1*, *CLGN*, *CRYZ*, *DPYSL5*, *RASSF3*, *NPTX2*, and *NEUROD4* showed the opposite expression pattern. Thus, variation in *CRTC1* expression within Group 3 β MB was associated with broader differences in tumor transcriptional profiles.

**Fig. 9.**
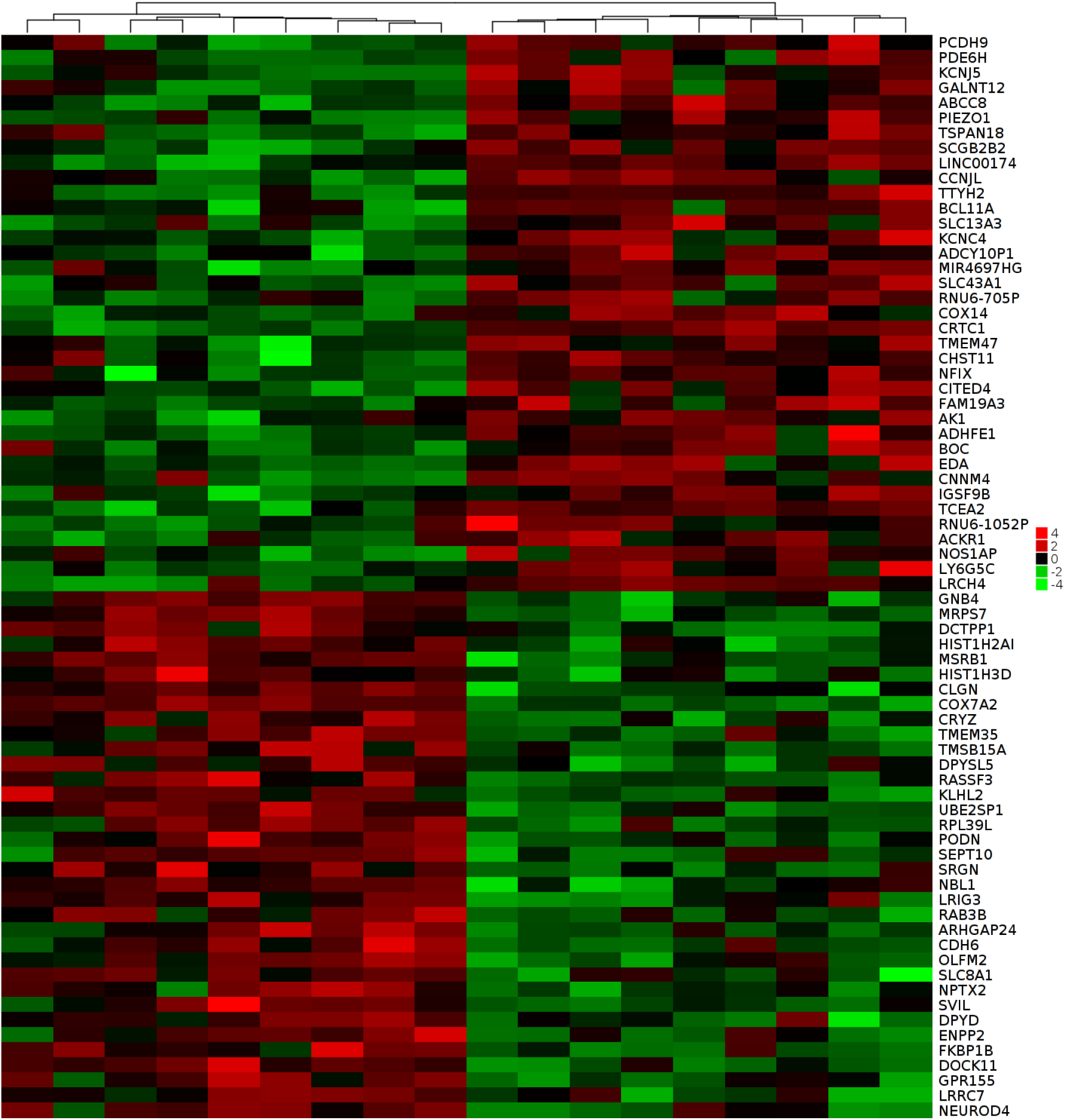
Differential gene expression (DGE) according to *CRTC1* expression in Group 3 β MB. Heatmap showing DEG between *CRTC1*-high and *CRTC1*-low Group 3 β tumors from the Cavalli cohort (GSE85217; *N* = 37) [3], generated using the GlioVis data portal (https://gliovis.bioinfo.cnio.es/) [29]. Rows represent genes and columns represent individual tumors, with hierarchical clustering illustrating differences in transcriptional profiles associated with *CRTC1* expression. Red indicates relatively higher expression and green indicates relatively lower expression.

### 3.7 *CRTC1* expression is associated with tumor microenvironment-related transcriptional features in Group 3 β MB

*CRTC1* expression in Group 3 β MB was inversely correlated with StromalScore (Spearman ρ = −0.325, *P* = 0.0500) and ESTIMATEScore (ρ = −0.336, *P* = 0.0421) and positively correlated with inferred tumor purity (ρ = 0.336, *P* = 0.0421), whereas its inverse correlation with ImmuneScore was weaker (ρ = −0.299, *P* = 0.0723). None of these rank correlations remained significant after correction across the four ESTIMATE-derived measures (FDR = 0.067 for StromalScore, ESTIMATEScore, and tumor purity; FDR = 0.072 for ImmuneScore). Complementary linear modeling within the Group 3 β subtype yielded significant inverse associations of *CRTC1* with StromalScore (β = −161.5 per 1-SD increase in *CRTC1*, *P* = 0.0104) and ESTIMATEScore (β = −288.2, *P* = 0.0142), together with a positive association with inferred tumor purity (β = 0.0105, *P* = 0.0142). These three associations remained significant after correction across the four measures (FDR = 0.0189), whereas the association with ImmuneScore did not (*P* = 0.0611) (Fig. 10).

**Fig. 10.**
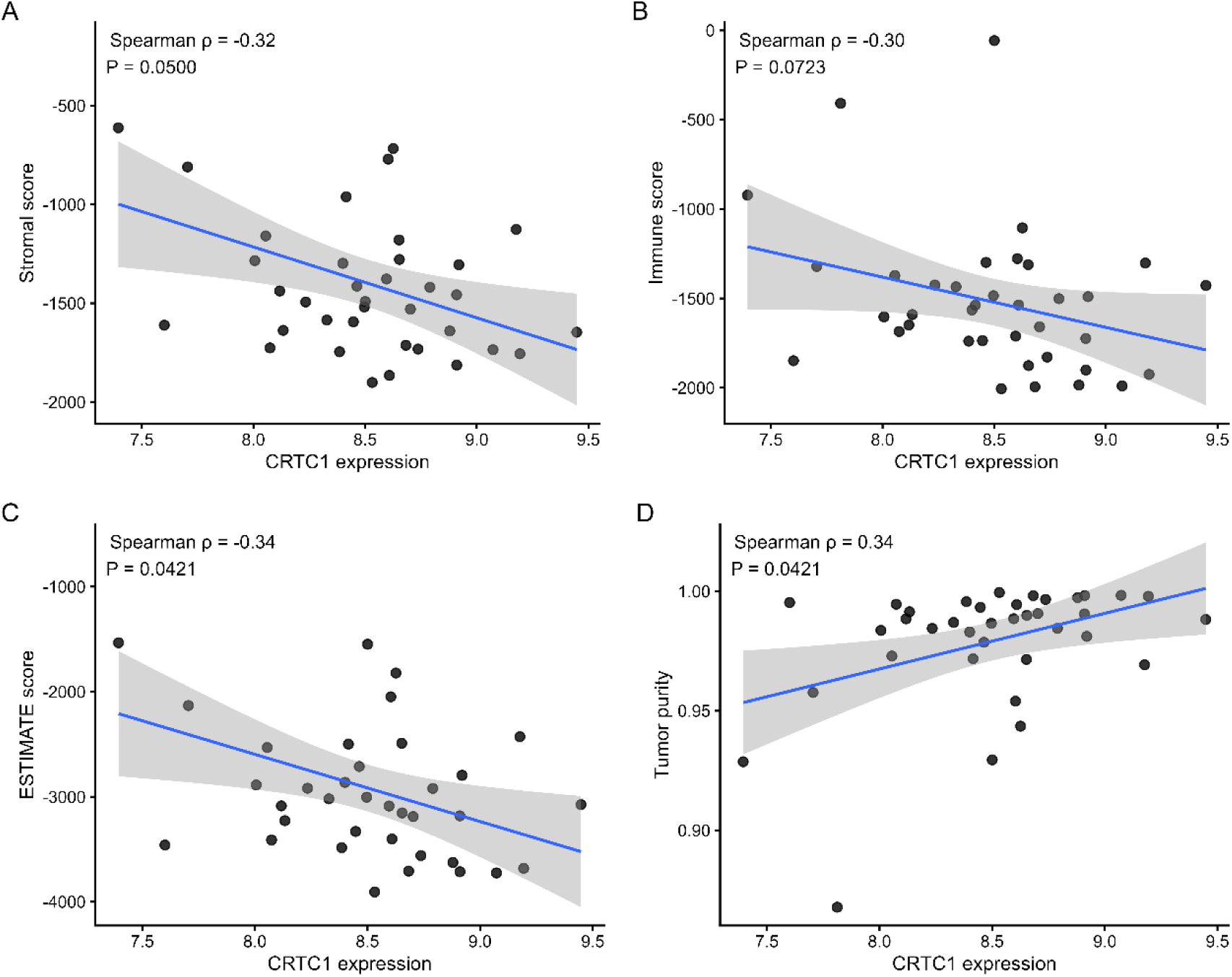
Association between *CRTC1* expression and tumor microenvironment estimates in Group 3 β MB. *CRTC1* expression was examined in relation to (**A**) StromalScore, (**B**) ImmuneScore, (**C**) ESTIMATEScore, and (**D)** inferred tumor purity in Group 3 β tumors from the Cavalli cohort (GSE85217; *N* = 37) [3]. Associations were evaluated using Spearman rank correlation. Lines indicate linear fits and shaded areas represent 95% CIs and are shown for visualization. *P* values shown in the panels are unadjusted. None of the four Spearman associations remained significant after BH correction (FDR = 0.067 for StromalScore, ESTIMATEScore, and tumor purity; FDR = 0.072 for ImmuneScore).

### 3.8 Specific cell-type enrichment signatures are associated with *CRTC1* within Group 3 β MB

We next examined xCell enrichment signatures specifically within Group 3 β tumors. Seven of the 64 cell-type signatures were nominally associated with *CRTC1* expression, of which two remained significant after correction across all 64 signatures. The strongest association was observed for the common myeloid progenitor (CMP) signature (ρ = 0.563, *P* = 2.88 × 10^−4^, FDR = 0.0184). *CRTC1* expression was also positively associated with the pericyte signature (ρ = 0.511, *P* = 0.00122, FDR = 0.0391) (Fig. 11). A smooth-muscle signature was inversely associated with *CRTC1* within Group 3 β (ρ = −0.476, *P*= 0.00290), although this association did not remain significant after correction across the 64 signatures (FDR = 0.0618). Other nominal Group 3 β associations included hepatocytes, CD4+ central-memory T cells, myocytes, and megakaryocytes, but none survived multiple-testing correction.

**Fig. 11.**
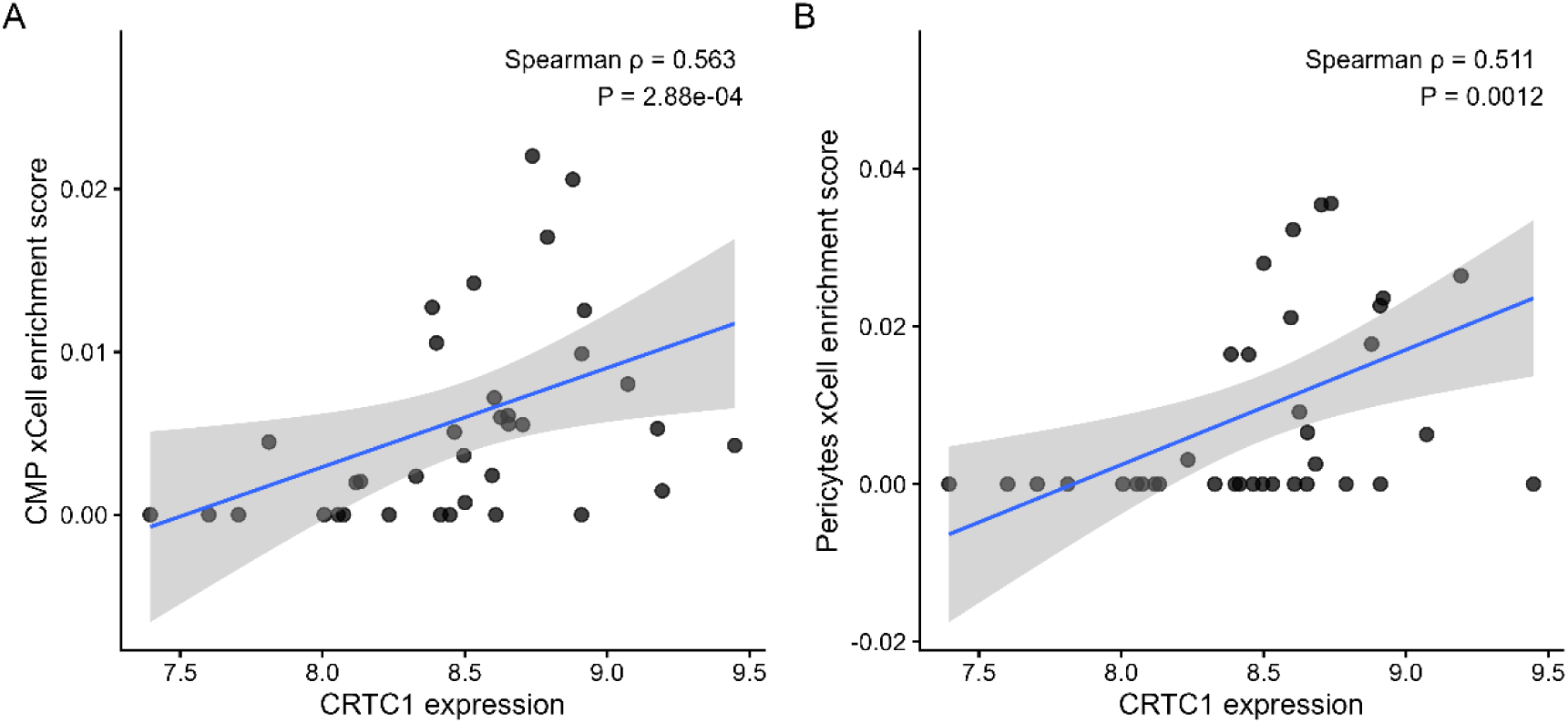
C*R*TC1 expression is associated with CMP- and pericyte-related enrichment signatures in Group 3 β MB. Associations between *CRTC1* expression and xCell enrichment scores for **A** CMPs and **B** pericytes were examined in Group 3 β tumors from the Cavalli cohort (GSE85217; *N* = 37) [3]. Spearman rank correlation showed positive associations between *CRTC1* expression and the CMP (ρ = 0.563, *P* = 2.88 × 10^−4^) and pericyte (ρ = 0.511, *P* = 0.00122) enrichment scores. These were the only two of 64 xCell cell-type signatures that remained significant after BH correction (CMP, FDR = 0.0184; pericytes, FDR = 0.0391). Lines indicate linear fits with 95% CIs and are shown for visualization. xCell scores represent transcriptional enrichment signatures and should not be interpreted as direct estimates of cell abundance.

In contrast to the specific CMP and pericyte signatures, none of the aggregate xCell measures was associated with *CRTC1* within Group 3 β: ImmuneScore (ρ = −0.039, *P* = 0.818), StromaScore (ρ = −0.051, *P* = 0.762), and MicroenvironmentScore (ρ = −0.048, *P* = 0.777). Thus, the Group 3 β xCell findings support associations with selected transcriptional cell-type signatures rather than a generalized shift in overall immune or stromal enrichment.

## 4 Discussion

Analysis of two independent developmental cerebellar datasets indicated that *CRTC1* expression is dynamically regulated within the human granule-cell lineage. In the Aldinger dataset, donor-level analyses showed lower expression in differentiated GNs than in RL cells, whereas expression in GCP remained similar to that in RL. The independent Sepp dataset revealed a different but complementary feature: a pronounced enrichment of *CRTC1* within the GC defined population, with substantially higher mean expression and a greater fraction of expressing nuclei than in GCP or either of the differentiating GC states. Together, these observations are more consistent with cell-state-specific developmental regulation of *CRTC1* than with uniform progressive induction or repression during granule-cell maturation. *CRTC1* was also expressed across all seven pediatric brain tumor classes but reached its highest median level in MB, with significantly higher expression than in every other tumor group examined. Together with the developmental cerebellar data, these findings are consistent with the possibility that high *CRTC1* expression in MB reflects retention or reactivation of transcriptional programs related to cerebellar development.

Biological mechanisms mediating neural plasticity can be amplified and hijacked in brain tumors [1, 2]. We have previously shown that expression of genes related to central nervous system development and synaptic transmission, including *GRIA1*-*GRIA4*, *GABRB1-GABRB3*, *TWIST1*, *NRP1*, and *PSEN1* can be associated with longer OS in MB patients in a tumor subgroup and subtype-dependent manner [24, 30–33]. Here we identify a subtype-restricted association between *CRTC1* expression and outcome in Group 3β MB. Group 3 MB tumors are associated with a poor prognosis, and it is essential to identify genes capable of improving risk stratification [34]. During human embryonal development, the RL generates all glutamatergic neurons within the cerebellum undergoing various morphological changes and exhibiting a complex proliferation pattern [35]. Group 3 MB present upregulated genes that define GNPs, which are proposed as cells of origin for this MB subgroup [7]. CREB, which is regulated by CRTC1, is highly expressed in its activated form in mature GNs [36] and low levels of active form of CREB are linked to a poor prognosis in Group 3 MB patients [37].

*MYC* amplification or overexpression is the primary molecular driver of Group 3 MB, which is also characterized by translocations of the *PVT1* oncogene, which encodes an lncRNA, with *MYC* and *NDRG*, and recurrent somatic copy number aberrations involving transforming growth factor β (TGF-β) signaling [38–40]. The transcription factor OTX2, which orchestrates a pro-tumorigenic splicing program mirrored in cerebellar RL development, maintains a stemness program in Group 3 MB [41]. An OTX2-associated regulatory network involving *PAX3* and *PAX6* controls Group 3 MB cell fate [42]. Within Group 3 MB, the Group 3 β subtype is particularly characterized by *OTX2* gain, *DDX31* loss, and *GFI1/1B* expression [3]. Our Group 3 β-specific coexpression and GO analyses showed functional enrichment concentrated among genes negatively correlated with *CRTC1*, which converged on mitochondrial organization and bioenergetics, including oxidative phosphorylation, respiratory electron transport, ATP synthesis, mitochondrial gene expression, and mitochondrial translation. This finding is particularly interesting in the context of the survival analysis, given that it suggests a low-*CRTC1* state characterized by increased expression of mitochondrial and bioenergetic programs within Group 3 β MB. The convergence of oxidative phosphorylation, respiratory-chain function, ATP synthesis, and mitochondrial gene expression suggests that the association extends beyond individual mitochondrial genes and reflects a broader transcriptional phenotype. Previous evidence linking CRTC1 with mitochondrial function in non-tumoral cells includes experiments showing that CRTC1 expression in primary muscle cells induces the endogenous mRNA of peroxisome proliferator-activated receptor gamma coactivator 1α (PGC-1α) and its downstream target genes in the mitochondrial respiratory chain and tricarboxylic acid (TCA) cycle in primary muscle cells. CRTC1 acts as one of the main upstream regulators of PGC-1α, which enhances mitochondrial oxidative capacity and mitochondrial gene expression [43]. The CRTC1/CREB pathway plays a role in regulating mitochondrial metabolism in microglia [44], and drives mitochondrial fragmentation and suppresses the effects of AMP-activated protein kinase (AMPK) on systemic mitochondrial metabolism in *C. elegans* [45]. In our analysis, genes positively correlated with *CRTC1* showed substantially more limited functional enrichment, primarily involving transcriptional elongation and neural developmental processes. Differential expression analysis comparing *CRTC1*-high and *CRTC1*-low tumors revealed distinct gene expression profiles. Genes distinguishing these states included regulators of transcription and differentiation as well as genes related to membrane signaling and cellular physiology. Together with the continuous coexpression analysis, these findings suggest that *CRTC1* expression marks distinct transcriptional states within Group 3 β MB, which may contribute to the subgroup-specific association between *CRTC1* and patient outcome.

The tumor microenvironment analyses indicated that *CRTC1* expression is associated with the transcriptional composition of Group 3 β MB. Higher CRTC1 expression was associated with lower stromal and overall ESTIMATE scores and correspondingly higher inferred tumor purity, with a weaker inverse trend for ImmuneScore. These findings initially suggest that *CRTC1*-high tumors may contain a reduced contribution from non-neoplastic microenvironment-associated transcription. None of the aggregate xCell immune, stromal, or microenvironment scores correlated with *CRTC1* within Group 3 β MB, whereas CMP- and pericyte-associated signatures remained significant after correction across all 64 cell-type signatures. However, xCell enrichment scores do not constitute direct measurements of cell abundance, and the presence of strong associations with signatures nominally assigned to hepatocytes, adipocytes, melanocytes, and other cell types not expected to constitute meaningful MB populations demonstrates the need for caution in assigning literal cellular identities to these signals. The CMP association should be interpreted as enrichment of a CMP-associated transcriptional signature rather than evidence for infiltration by bona fide common myeloid progenitors. Validation using single-cell or spatial transcriptomic data would be required to determine which cellular compartments contribute to this signal. These microenvironment findings also provide an additional dimension to the Group 3 β MB survival association with *CRTC1*, but they do not demonstrate that microenvironmental differences mediate the relationship between *CRTC1* and patient survival. At present, the data support a model in which *CRTC1* expression marks a distinct transcriptional state in Group 3 β MB that encompasses both tumor-intrinsic and microenvironment-associated features.

This study has several limitations. Because the analyses are based on bulk tumor transcriptomes and observational associations, they cannot establish whether CRTC1 directly regulates the differentially expressed genes or instead reflects underlying differences in tumor cell state or cellular composition. Also, the survival analysis must be interpreted as purely associative and was restricted to the 27 Group 3 β tumors with evaluable outcome data. Moreover, transcript abundance does not directly establish mitochondrial respiratory activity, and the present analyses cannot determine whether CRTC1 directly regulates mitochondrial programs or whether both features reflect a broader biological state of Group 3 β tumor cells. Computational genomic studies interrogating established datasets such as the present one should be considered exploratory and hypothesis-generating. Functional experiments will therefore be required to establish whether modulation of CRTC1 alters mitochondrial metabolism and whether such changes contribute to tumor behavior.

## 5 Conclusion

In summary, our findings identify *CRTC1* as a developmentally regulated transcriptional coactivator gene with heterogeneous expression across MB molecular groups and subtypes and a particularly strong clinical association within Group 3 β MB. High *CRTC1* expression was associated with longer patient survival, whereas low *CRTC1* expression was linked to enrichment of mitochondrial and bioenergetic transcriptional programs. Together with the distinct transcriptional profiles and selected cell-type-associated signatures observed according to *CRTC1* expression, these results indicate that *CRTC1* marks biologically distinct states within Group 3 β MB. Experimental studies will be required to determine whether CRTC1 contributes functionally to these states and whether its subtype-specific association with outcome has prognostic or therapeutic relevance.

## Author Contributions

Conceptualization, J.V., R.R.; methodology, J.V., R.R.; formal analysis, J.V., R.R.; investigation, J.V., M.M.L., A.T.B., C.B.F., M.J., R.R., data curation, J.V., R.R.; writing – original draft, J.V., R.R.; Writing – review & editing, J.V., M.M.L., A.T.B., C.B.F., M.J., R.R.; Supervision, R.R.; Project administration, J.V.,A.T.B., C.B.F., M.J., R.R.; Funding acquisition, J.V., A.T.B., C.B.F., M.J., R.R.

## Data Availability

All datasets analyzed in this study are publicly available. Gene expression data from human brain areas were obtained from The Human Protein Atlas (HPA; https://www.proteinatlas.org/ENSG00000105662-CRTC1/brain). Gene expression data for the human cerebellum across development were obtained from the datasets described by Aldinger et al. [25] and Sepp et al. [26]. Gene expression data in pediatric brain tumors were obtained from the Open Pediatric Brain Tumor Atlas (OpenPBTA) [27] (https://github.com/AlexsLemonade/OpenPBTA-analysis). Gene expression and patient survival data in MB were obtained the Cavalli MB dataset (GSE85217) [3]. Single-cell RNA sequencing (scRNA-seq) data were obtained from dataset GSE156053 [28] and explored using the Pediatric Neuro-oncology Cell Atlas interactive platform (https://www.pneuroonccellatlas.org/). Differential gene expression was analyzed using the GlioVis data portal (https://gliovis.bioinfo.cnio.es/). R scripts required to reproduce gene expression analysis in MB are publicly available at the GitHub repository archived as https://github.com/rafaelroesler/CRTC1-medulloblastoma.

## Declarations

### Consent to Participate

Not applicable.

### Consent to Publish

Not applicable.

### Human Ethics and Consent to Participate

This study was based exclusively on analyses of publicly available, de-identified transcriptomic datasets and publicly accessible online bioinformatic resources. No human participants were recruited, no human tissue or biological specimens were collected or analyzed, no identifiable patient information was accessed, and no animal experiments were performed. Therefore, institutional ethics committee approval and informed consent were not required.

## References

1. Monje M. The neuroscience of brain cancers. Neuron. 2025;113(17):2734–9. doi: 10.1016/j.neuron.2025.07.012

2. Monje M. The neuroscience of childhood brain cancers. Sci Transl Med. 2026;18(860):eaeg9138. doi: 10.1126/scitranslmed.aeg9138

3. Cavalli FMG, Remke M, Rampasek L, Peacock J, Shih DJH, Luu B, et al. Intertumoral heterogeneity within medulloblastoma subgroups. Cancer Cell. 2017;31(6):737–54.e6. doi: 10.1016/j.ccell.2017.05.005

4. Gibson P, Tong Y, Robinson G, Thompson MC, Currle DS, Eden C, et al. Subtypes of medulloblastoma have distinct developmental origins. Nature. 2010;468(7327):1095–9. doi: 10.1038/nature09587

5. Vladoiu MC, El-Hamamy I, Donovan LK, Farooq H, Holgado BL, Sundaravadanam Y, et al. Childhood cerebellar tumours mirror conserved fetal transcriptional programs. Nature. 2019;572(7767):67–73. doi: 10.1038/s41586-019-1158-7

6. Hovestadt V, Smith KS, Bihannic L, Filbin MG, Shaw ML, Baumgartner A, et al. Resolving medulloblastoma cellular architecture by single-cell genomics. Nature. 2019;572(7767):74–9. doi: 10.1038/s41586-019-1434-6

7. Smith KS, Bihannic L, Gudenas BL, Haldipur P, Tao R, Gao Q, et al. Unified rhombic lip origins of group 3 and group 4 medulloblastoma. Nature. 2022;609(7929):1012–20. doi: 10.1038/s41586-022-05208-9

8. Northcott PA, Buchhalter I, Morrissy AS, Hovestadt V, Weischenfeldt J, Ehrenberger T, et al. The whole-genome landscape of medulloblastoma subtypes. Nature. 2017;547(7663):311–7. doi: 10.1038/nature22973

9. Northcott PA, Korshunov A, Pfister SM, Taylor MD. The clinical implications of medulloblastoma subgroups. Nat Rev Neurol. 2012;8(6):340–51. doi: 10.1038/nrneurol.2012.78

10. Kovács KA, Steullet P, Steinmann M, Do KQ, Magistretti PJ, Halfon O, et al. TORC1 is a calcium- and cAMP-sensitive coincidence detector involved in hippocampal long-term synaptic plasticity. Proc Natl Acad Sci U S A. 2007;104(11):4700–5. doi: 10.1073/pnas.0607524104

11. Zhou Y, Wu H, Li S, Chen Q, Cheng XW, Zheng J, et al. Requirement of TORC1 for late-phase long-term potentiation in the hippocampus. PLoS One. 2006;1(1):e16. doi: 10.1371/journal.pone.0000016

12. Ch’ng TH, Uzgil B, Lin P, Avliyakulov NK, O’Dell TJ, Martin KC. Activity-dependent transport of the transcriptional coactivator CRTC1 from synapse to nucleus. Cell. 2012;150(1):207–21. doi: 10.1016/j.cell.2012.05.027

13. Tan BZ, Cuevas JNS, Yoshida R, Uematsu A, Koivumaa J, Ishizu Y, et al. A neuromodulatory circuit-to-molecular pathway for reformatting aversive memories during recall. Neuron. 2026;114(10):1848–62.e7. doi: 10.1016/j.neuron.2026.01.006

14. Nonaka M, Kim R, Fukushima H, Sasaki K, Suzuki K, Okamura M, et al. Region-specific activation of CRTC1-CREB signaling mediates long-term fear memory. Neuron. 2014;84(1):92–106. doi: 10.1016/j.neuron.2014.08.049

15. España J, Valero J, Miñano-Molina AJ, Masgrau R, Martín E, Guardia-Laguarta C, et al. beta-Amyloid disrupts activity-dependent gene transcription required for memory through the CREB coactivator CRTC1. J Neurosci. 2010;30(28):9402–10. doi: 10.1523/JNEUROSCI.2154-10.2010

16. Wilson EN, Abela AR, Do Carmo S, Allard S, Marks AR, Welikovitch LA, et al. Intraneuronal amyloid beta accumulation disrupts hippocampal CRTC1-dependent gene expression and cognitive function in a rat model of Alzheimer disease. Cereb Cortex. 2017;27(2):1501–11. doi: 10.1093/cercor/bhv332

17. Yan P, Xue Z, Li D, Ni S, Wang C, Jin X, et al. Dysregulated CRTC1-BDNF signaling pathway in the hippocampus contributes to Aβ oligomer-induced long-term synaptic plasticity and memory impairment. Exp Neurol. 2021;345:113812. doi: 10.1016/j.expneurol.2021.113812

18. Parra-Damas A, Valero J, Chen M, España J, Martín E, Ferrer I, et al. Crtc1 activates a transcriptional program deregulated at early Alzheimer’s disease-related stages. J Neurosci. 2014;34(17):5776–87. doi: 10.1523/JNEUROSCI.5288-13.2014

19. Amelio AL, Fallahi M, Schaub FX, Zhang M, Lawani MB, Alperstein AS, et al. CRTC1/MAML2 gain-of-function interactions with MYC create a gene signature predictive of cancers with CREB-MYC involvement. Proc Natl Acad Sci U S A. 2014;111(32):E3260–8. doi: 10.1073/pnas.1319176111

20. Chen Z, Chen J, Gu Y, Hu C, Li JL, Lin S, et al. Aberrantly activated AREG-EGFR signaling is required for the growth and survival of CRTC1-MAML2 fusion-positive mucoepidermoid carcinoma cells. Oncogene. 2014;33(29):3869–77. doi: 10.1038/onc.2013.348

21. Feng X, Shi Y, Yuan F, Hu Y, Tang X, Zhang W, et al. CRTC1 enhances PD-L1-mediated tumor immunosuppression in non-small cell lung cancer via the Notch1/Akt signaling pathway. Front Immunol. 2025;16:1658679. doi: 10.3389/fimmu.2025.1658679

22. Komiya T, Coxon A, Park Y, Chen WD, Zajac-Kaye M, Meltzer P, et al. Enhanced activity of the CREB co-activator Crtc1 in LKB1 null lung cancer. Oncogene. 2010;29(11):1672–80. doi: 10.1038/onc.2009.453

23. Schumacher Y, Aparicio T, Ourabah S, Baraille F, Martin A, Wind P, et al. Dysregulated CRTC1 activity is a novel component of PGE2 signaling that contributes to colon cancer growth. Oncogene. 2016;35(20):2602–14. doi: 10.1038/onc.2015.283

24. Vanini J, Thomaz A, Lupatini MM, Brunetto AT, de Farias CB, Jaeger M, et al. PSEN1 expression identifies a developmentally distinct favourable-prognosis state in SHH α medulloblastoma. Int J Dev Neurosci. 2026;86(6):e70178. doi: 10.1002/jdn.70178

25. Aldinger KA, Thomson Z, Phelps IG, Haldipur P, Deng M, Timms AE, et al. Spatial and cell type transcriptional landscape of human cerebellar development. Nat Neurosci. 2021;24(8):1163–75. doi: 10.1038/s41593-021-00872-y

26. Sepp M, Leiss K, Murat F, Okonechnikov K, Joshi P, Leushkin E, et al. Cellular development and evolution of the mammalian cerebellum. Nature. 2024;625(7996):788–96. doi: 10.1038/s41586-023-06884-x

27. Shapiro JA, Gaonkar KS, Spielman SJ, Savonen CL, Bethell CJ, Jin R, et al. Children’s Brain Tumor Network; Pacific Pediatric Neuro-Oncology Consortium. OpenPBTA: The Open Pediatric Brain Tumor Atlas. Cell Genom. 2023;3(7):100340. doi: 10.1016/j.xgen.2023.100340

28. Riemondy KA, Venkataraman S, Willard N, Nellan A, Sanford B, Griesinger AM, et al. Neoplastic and immune single-cell transcriptomics define subgroup-specific intra-tumoral heterogeneity of childhood medulloblastoma. Neuro Oncol. 2022;24(2):273–86. doi: 10.1093/neuonc/noab135

29. Bowman RL, Wang Q, Carro A, Verhaak RG, Squatrito M. GlioVis data portal for visualization and analysis of brain tumor expression datasets. Neuro Oncol. 2017;19(1):139–41. doi: 10.1093/neuonc/now247

30. Coelho KMPA, Dalmolin M, Pedack FR, Pickler JG, Barbosa C, Recalde J, et al. Opposite patterns of association of TWIST1 expression with patient survival in SHH and Group 4 medulloblastoma. Childs Nerv Syst. 2025;41(1):277. doi: 10.1007/s00381-025-06913-2

31. de Araújo MA, Malafaia O, Ribas Filho JM, Fratini L, Roesler R, Isolan GR. Low expression of the NRP1 gene is associated with shorter overall survival in patients with sonic hedgehog and Group 3 medulloblastoma. Int J Mol Sci. 2023;24(14):11601. doi: 10.3390/ijms241411601

32. Monteiro JM, Dalmolin M, Fernandes MAC, Ramos JIR, Ribas CAPM, Tabushi FI, et al. High expression of GABA_A_ receptor β subunit genes is associated with longer overall survival in medulloblastoma. Brain Sci. 2024;14(11):1146. doi: 10.3390/brainsci14111146

33. Saciloto B, Dalmolin M, Marcolin JC, Lichtenfels M, Vanini J, Roesler IBS, et al. Subgroup-specific associations of GRIA genes encoding AMPA glutamate receptor subunits with patient survival in medulloblastoma. Neuromolecular Med. 2026;28(1):36. doi: 10.1007/s12017-026-08935-8

34. Taylor MD, Northcott PA, Korshunov A, Remke M, Cho YJ, Clifford SC, et al. Molecular subgroups of medulloblastoma: the current consensus. Acta Neuropathol. 2012;123(4):465–72. doi: 10.1007/s00401-011-0922-z

35. Haldipur P, Aldinger KA, Bernardo S, Deng M, Timms AE, Overman LM, et al. Spatiotemporal expansion of primary progenitor zones in the developing human cerebellum. Science. 2019;366(6464):454–60. doi: 10.1126/science.aax7526

36. Dworkin S, Mantamadiotis T. Targeting CREB signalling in neurogenesis. Expert Opin Ther Targets. 2010;14(8):869–79. doi: 10.1517/14728222.2010.501332

37. Armandari I, Zomerman WW, Plasschaert SLA, Smit MJ, Martini TEI, de Camargo Magalhães ES, et al. CREB signaling activity correlates with differentiation and survival in medulloblastoma. Sci Rep. 2021;11(1):16077. doi: 10.1038/s41598-021-95381-0

38. Katsushima K, Joshi K, Yuan M, Romero B, Batish M, Stapleton S, et al. A therapeutically targetable positive feedback loop between lnc-HLX-2-7, HLX, and MYC that promotes group 3 medulloblastoma. Cell Rep. 2024;43(3):113938. doi: 10.1016/j.celrep.2024.113938

39. Northcott PA, Shih DJ, Peacock J, Garzia L, Morrissy AS, Zichner T, et al. Subgroup-specific structural variation across 1,000 medulloblastoma genomes. Nature. 2012;488(7409):49–56. doi: 10.1038/nature11327

40. Roussel MF, Robinson GW. Role of MYC in medulloblastoma. Cold Spring Harb Perspect Med. 2013;3(11):a014308. doi: 10.1101/cshperspect.a014308.

41. Saulnier O, Zagozewski J, Liang L, Hendrikse LD, Layug P, Gordon V, et al. A group 3 medulloblastoma stem cell program is maintained by OTX2-mediated alternative splicing. Nat Cell Biol. 2024;26(8):1233–46. doi: 10.1038/s41556-024-01460-5

42. Zagozewski J, Shahriary GM, Morrison LC, Saulnier O, Stromecki M, Fresnoza A, et al. An OTX2-PAX3 signaling axis regulates Group 3 medulloblastoma cell fate. Nat Commun. 2020;11(1):3627. doi: 10.1038/s41467-020-17357-4

43. Wu Z, Huang X, Feng Y, Handschin C, Feng Y, Gullicksen PS, et al. Transducer of regulated CREB-binding proteins (TORCs) induce PGC-1alpha transcription and mitochondrial biogenesis in muscle cells. Proc Natl Acad Sci U S A. 2006;103(39):14379–84. doi: 10.1073/pnas.0606714103

44. Fang N, Wang Y, Chen Y, Wang Y, Xu J, Xie Y, et al. SIK2 mediated mitochondrial homeostasis in spinal cord injury: modulating oxidative stress and the AIM2 inflammasome via CRTC1/CREB signaling. J Neuroinflammation. 2025;22(1):283. doi: 10.1186/s12974-025-03606-0

45. Burkewitz K, Morantte I, Weir HJM, Yeo R, Zhang Y, Huynh FK, et al. Neuronal CRTC-1 governs systemic mitochondrial metabolism and lifespan via a catecholamine signal. Cell. 2015;160(5):842–55. doi: 10.1016/j.cell.2015.02.004

